# dingo: a Python package for metabolic flux sampling

**DOI:** 10.1101/2023.06.18.545486

**Authors:** Apostolos Chalkis, Vissarion Fisikopoulos, Elias Tsigaridas, Haris Zafeiropoulos

## Abstract

**Summary:** We present `dingo`, a Python package that supports a variety of methods to sample from the flux space of metabolic models, based on state-of-the-art random walks and rounding methods. For uniform sampling `dingo`’s implementation of the Multiphase Monte Carlo Sampling algorithm, provides a significant speed-up and outperforms existing software. Indicatively, `dingo` can sample from the flux space of the largest metabolic model up to now (Recon3D) in less than 30 hours using a personal computer, under several statistical guarantees; this computation is out of reach for other similar software. In addition, supports common analysis methods, such as Flux Balance Analysis (FBA) and Flux Variability Analysis (FVA), and visualization components. `dingo` contributes to the arsenal of tools in metabolic modeling by enabling flux sampling in high dimensions (in the order of thousands).

**Availability and implementation:** https://github.com/GeomScale/dingo

## 1 Introduction

Metabolic models enhance the study of the relationship between genotype and phenotype in an attempt to elucidate the mechanisms that govern the physiology and the growth of a species and/or a community (Morris *et al*., 2020). By optimizing a linear objective function over a polytope, Flux Balance Analysis (FBA) identifies a single optimal flux distribution (Orth *et al*., 2010). Flux Variability Analysis (FVA) reveals the limits of the solution space (Gudmundsson and Thiele, 2010). Contrary to FBA and FVA, flux sampling is an unbiased method, as it does not depend on the selection of the objective function. It allows us to cover all the possible flux values by estimating a probability distribution for the flux value of a certain reaction (Schellenberger and Palsson, 2009).

The ability to sample (efficiently) points from the convex polytope corresponding to a metabolic model allows us to investigate its whole solution space. This way, we can obtain a more detailed insight of a system at steady state; where the production rate of each metabolite equals its consumption rate. Alternatively, we can perform flux sampling after optimizing an objective function, and approximate the flux distributions in optimal scenarios.

Even though flux sampling has proved itself by delivering great insights in a range of applications (Herrmann *et al*., 2019), high dimensionality- and anisotropy-oriented limitations (Schellenberger and Palsson, 2009) force the current implementations to struggle or even to fail in several cases (Fallahi *et al*., 2020). A range of Markov Chain Monte Carlo (MCMC) algorithms and implementations have been developed to address this obstacle (Fallahi *et al*., 2020) (see Supplementary File - Section 1). In this setting, we present `dingo`, a Python package that supports efficient flux sampling, based on a variety of state-of-the-art MCMC sampling algorithms; it also provides classical FBA and FVA methods and advanced visualisations.

## 2 Implementation

`dingo` is an open-source Python package that exploits the efficiency of `volesti`, an open-source `C++` software library that implements high-dimensional MCMC sampling and volume approximation algorithms.

`dingo` supports a variety of MCMC algorithms for uniform sampling. Among them, the Multiphase Monte Carlo Sampling (MMCS) algorithm (Chalkis *et al*., 2021) has been reported as the most efficient algorithm in practice. MMCS unifies rounding and sampling of a convex polytope in one pass, obtaining both upon termination.

In addition to the Python interface, `dingo` enables the performance of the MMCS algorithm in parallel threads and uses the state-of-the-art linear programming solvers of `Gurobi` Gurobi Optimization, LLC (2023). It ensures the quality of the output samples using two widely used diagnostics, the Effective Sample Size (ESS) (Geyer, 1992) and the Potential Scale Reduction Factor (PSRF) (Gelman and Rubin, 1992); `dingo` guarantees bounded values for both diagnostics for the returned sample. In addition to the MMCS algorithm, it also supports the Random Directions Hit-and-Run (RDHR) (Smith, 1984), the Coordinate Directions Hit-and-Run (CDHR) (Smith, 1984), the Dikin (Kannan and Narayanan, 2012), the John and Vaidya (Chen *et al*., 2018) and the Ball Walk (Lovász *et al*., 1997) sampling algorithms.

We ensure the correctness of `dingo`’s functionality using a set of unit tests running on a continuous integration platform. `dingo` supports the three main formats for metabolic models (.xml, .json and .mat). A tutorial is available as a Google Colab notebook ^1^.

## 3 Performance comparison and illustrations

Currently, the most efficient way to perform flux sampling, to the best of our knowledge, is to combine the `PolyRound` Python package (Theorell *et al*., 2021) (for rounding the polytope) with CDHR sampling algorithm as implemented in the the `HOPS` C++ library (Jadebeck *et al*., 2021). We compare `dingo` against the combination of `PolyRound` and `hopsy` (the Python interface of `HOPS`), over a set of models having dimension from *∼* 100 to *>* 13, 000. In all cases, we perform a pre-processing step using `PolyRound`. As for `PolyRound`/`hopsy`, the polytopes are rounded with `PolyRound` and sampled with `hopsy`. On the other hand, `dingo` performs rounding and sampling in one step using the MMCS algorithm (see also Supplementary File - Section 2). To ensure that the quality of the sample provides an accurate approximation of the target distribution, we require an ESS of 1, 000 and a PSRF of at most 1.1, in all cases. To our knowledge `dingo` is the only software that provides this combined statistical guarantee. For all the tested models `dingo` is faster than `PolyRound` / `hopsy`. Moreover, `dingo`’s added value highlights as the model’s dimensions increases (see Fig. 1). Indicatively, `dingo` is able to sample the latest version of the human metabolic network, the Recon3D model (Brunk *et al*., 2018), in *∼* 1 day, using modest hardware; while after 10 days, `hopsy` did not converge. Notably, `dingo`’s run-time (for all the models but one) is comparable with the time that `PolyRound` requires to perform solely the rounding step (Supplementary Fig. 1).

**Fig. 1.**
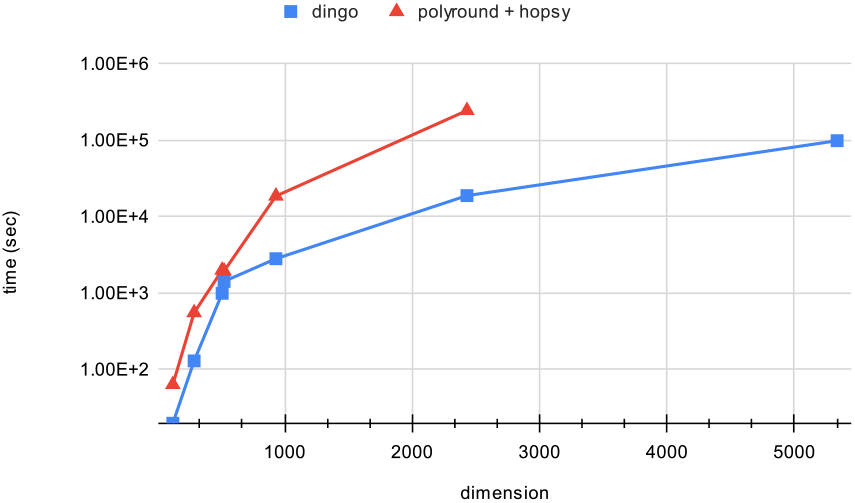
`dingo` vs. `PolyRound`/`hopsy` computing times when sampling from the flux space of 7 GEMs corresponding to polytopes of dimension ranging from 122 to 5335, under the same statistical guarantees. PolyRound is used for rounding and hopsy for sampling from the rounded polytope. dingo’s run-time corresponds to both rounding and sampling, starting from the non-rounded polytope (i.e. same as the input of PolyRound).

To demonstrate `dingo`’s flux sampling and illustrations tools in a real-world scenario, we use the integrated human alveolar macrophage model with the virus biomass objective function (VBOF) of Sars-Cov-2 (Renz *et al*., 2020) (see Supplementary File - Section 3). Notably, our findings confirm the authors’ indicating Guanylate Kinase 1 as a potential therapeutic target.

## 4 Conclusions

`dingo` is a Python package that employs efficient C++ MCMC implementations from `volesti` library. It provides a novel, efficient implementation of the MMCS algorithm that offers rounding and sampling in one pass. It supports a variety of MCMC algorithms and classical methods as FBA and FVA. `dingo` outperforms the existing software for sampling the flux space. It also offers statistical and illustration tools, like copula estimation, that can help the user to extract useful information about the model (see Supplementary File – Section 3). `dingo` facilitates the survey of the largest models available for the time being assuring for the first time high quality of the samples returned. Moreover, it requires minimum computational resources requirements and aims to support a broad spectrum of research and application needs via a user-friendly design.

## Acknowledgements

We thank Area 52 lab in University College Dublin for allowing us to use the *bob* server to run the comparison experiments.

## Footnotes

1 https://colab.research.google.com/github/GeomScale/dingo/blob/develop/tutorials/dingo_tutorial.ipynb.

