## Supplementary Information for "dingo: a Python package for metabolic flux sampling"

### `dingo`: a Python package for metabolic flux sampling Supplementary material

#### 1 Convex polytope sampling algorithms and available implementations

Uniform sampling from the flux space of high dimensional polytopes is challenging from a computational point-of-view. It is an active area in computational geometry and statistics, and thus, several sampling algorithms have been developed over the years and are consequently implemented in various software packages (see Table 1).

A big number of algorithms and their corresponding implementations exploit Markov Chain Monte Carlo (MCMC) methods, as MCMC sampling is considered the most efficient choice for high dimensional problems. In this setting, the widely used hit-and-run family of walks generates a chain of points by taking steps of random length inside the polytope in randomly generated directions. Each variant in this family generates the set of possible directions in a different way, e.g., Coordinate Hit-and-Run, in each step, picks randomly a line parallel to the axis and then a random point on the part of the line that lies inside the polytope. Another family of walks are the affine-invariant ones, namely the Dikin [8], Vaidya and John Walk [2]. These walks select the new point from the interior of an inscribed to the polytope ellipsoid. Each variant in this family defines that ellipsoid in a different way. The GAPSPLIT algorithm tries to divide the solution space uniformly to calculate random samples. To support flux sampling on the solution space of metabolic models, several software packages exists that implement these algorithms and also provide interfaces for a variety of languages, such as MATLAB (`cobra` [4, 5], `gapsplit` [9], optGpSampler [12]), Python (`gapsplit`, `optGpSampler`, `HOPS` [6]), C++ (`HOPS`, `polytopewalk` [2] [1]). The most efficient algorithm today is reported to be the Coordinate Hit-and-Run with Rounding (CHRR), while it has been the algorithm of choice for the state-of-the-art software (see `HOPS` and `cobra`). CHRR, before sampling, applies a pre-processing step, called rounding, so that the efficiency of the random walk in the sampling phase is improved. For sampling it employs the Coordinate Hit-and-Run algorithm.

Typically, rounding is applied on the polytope before sampling so that its roundness is improved. Roughly speaking, there are two notions of roundness: a polytope in John position [7] and a well-rounded polytope [11]. Both contain the unit ball, while the first is contained in a ball of radius $O^*(n)$ and the second in a ball of radius $O^*(\sqrt{n})$. We know that a polytope in isotropic position [14] is also well-rounded. Despite the fact that John position, in the worst case, is worse than well roundness, it is the main choice in the existing software for rounding because it is computationally easier to achieve. The Python package `PolyRound` seems to provide the most efficient implementation to achieve John position by computing the largest inscribed ellipsoid in the polytope and apply to it the transformation that maps the ellipsoid to the unit ball (`cobra` and `HOPS` also implement this rounding method).

`dingo` provides an efficient implementation of the Multiphase Monte Carlo Sampling (MMCS) algorithm by Chalkis *et al.* [1]. MMCS constructs a sequence of polytopes (phases) such that sampling is accelerated in each phase. All the samples are mapped back to the initial polytope, while both a rounded polytope and a sample set are obtained upon termination. In each phase,

[1] `https://github.com/yuachen/polytopewalk`

| Sampling algorithm | Software | | | | | |
|---|---|---|---|---|---|---|
| | cobra [4, 5] | HOPS [6] | polytopewalk [2] | gapsplit [9] | optGpSampler [12] | dingo |
| GAPSPLIT [9] | | | | x | | |
| Random Directions Hit-and-Run [19] | x | x | | | | x |
| Artificial Centering Hit-and-Run [13, 5] | x | | | | x | |
| Coordinate Hit-and-Run [?] | x | x | | | | x |
| Dikin Walk [8] | | x | x | | | x |
| Vaidya [2] | | | x | | | x |
| John Walk [2] | | | x | | | x |
| Ball Walk [10] | | | x | | | x |
| Billiard Walk [3] | | | | | | x |
| CHRR* [4] | x | x | | | | x |
| Multiphase Monte Carlo Sampling* [1] | | | | | | x |

Table 1: Selection of convex polytope sampling algorithms and related software packages. We denote with asterisk (*) the algorithms that also perform rounding to the polytope.

MMCS samples from the corresponding polytope and then applies to it the same transformation that maps the sample to an isotropic position, to obtain the polytope of the next phase. For sampling it employs an efficient version of the Billiard Walk algorithm [1]. MMCS computes the total Effective Sample Size (ESS) and the overall Potential Scale Reduction Factor (PSRF) through the phases and stops when both achieve a certain threshold given by the user. Finally, MMCS checks if the transformed polytope is in isotropic position before ESS and PSRF exceed their targets and stops applying transformations to further round the polytope. MMCS is the first method that unifies rounding and sampling in one pass and it is typically applied on a non-rounded polytope. Thus, the rounding preprocess is not necessary for MMCS. As shown in [1], MMCS outperforms CHRR as implemented in the `cobra` package by one or two orders of magnitude, depending on the dimension. `dingo` provides an optimized C++ implementation of MMCS accessible from a Python interface. On top of that, it provides a parallel implementation of MMCS, and for completeness, it provides a variety of MCMC sampling and rounding algorithms—including the aforementioned rounding method that brings the polytope to John position and the CHRR sampling algorithm.

| | | PolyRound | hopsy | | | dingo | |
|---|---|---|---|---|---|---|---|
| **Model name** | **d** | **rounding** | **sampling** | **total** | **PSRF** | **MMCS** | **PSRF** |
| iJN746 | 122 | 32.09 | 31.58 | 63.67 | 1.0062 | 19.81 | 1.0043 |
| iAT_PLT_636 | 289 | 132.43 | 421.96 | 554.39 | 1.0078 | 129.87 | 1.0065 |
| iSDY_1059 | 509 | 289.80 | 1707.20 | 1997.00 | 1.0100 | 993.32 | 1.0119 |
| iAF1260 | 524 | 290.17 | 1651.17 | 1941.34 | 1.0074 | 1422.20 | 1.0021 |
| recon1 | 931 | 2068.21 | 16535.48 | 18603.69 | 1.0089 | 2817.65 | 1.0034 |
| recon2d | 2430 | 14245.17 | 230636.15 | 244881.32 | 1.0100 | 18855.43 | 1.0870 |
| recon3d (latest) | 5335 | 55525.24 | NA | NA | NA | 98620.30 | 1.0040 |

Table 2: Time (in seconds) performance for sampling a metabolic model with `hopsy` (rounding with `PolyRound`) and `dingo` packages. The dimension $d$ is after preprocessing. For `hopsy` we use a thinning of $100d$ for all models but Recon3D where we use $200d$ as suggested by the authors [6]. NA means that after 10 days, `hopsy` was not able to converge and the process stopped at ESS=430, PSRF=1.024.

#### 2 Performance comparisons

To illustrate the efficiency of `dingo` we compare its MMCS implementation against the best existing implementation of CHRR[2]. `HOPS` library is reported to be six times faster for sampling than `cobra` [6] while both implementing CHRR. However, since `PolyRound` provides the most

[2]The evaluation of the efficiency of the CHRR implementation in `dingo` is out of the scope of this paper.

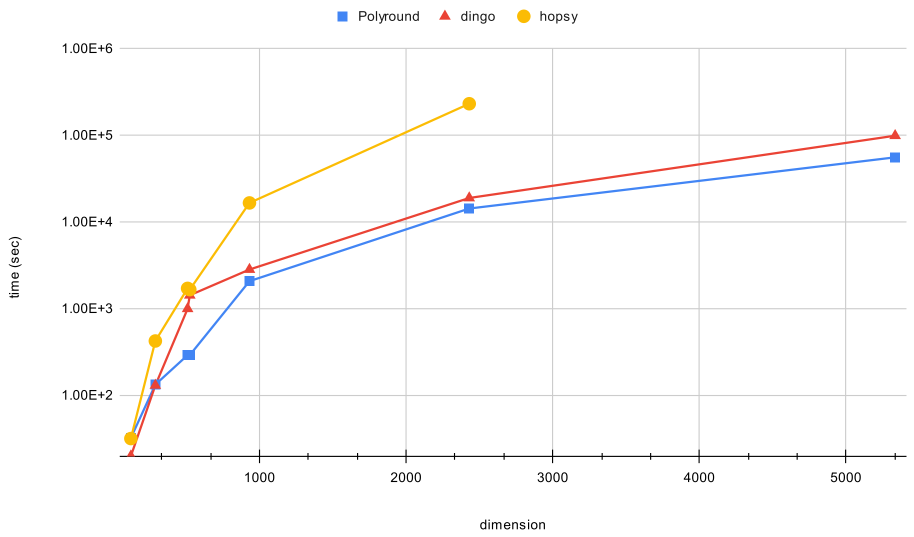


Figure 1: `dingo` (both rounding and sampling) vs. `PolyRound` (for rounding) and `hopsy` (for sampling) computing times when sampling from the flux space of 7 GEMs corresponding to polytopes of dimension ranging from 122 to 5335, under the same statistical guarantees.

efficient implementation for the rounding step in CHRR, we compare against the combination of `PolyRound` and `HOPS`. In particular, we compute the rounded polytope with `PolyRound` and we sample from the rounded polytope using `hopsy`, the python API of `HOPS`[3]. In `dingo` we use the non-rounded polytope. To obtain the polytope corresponding to a metabolic network, we use the routines of `PolyRound` as it also provides efficient facet redundancy removal implementations. For an overview of the workflow we refer to Figure 2.

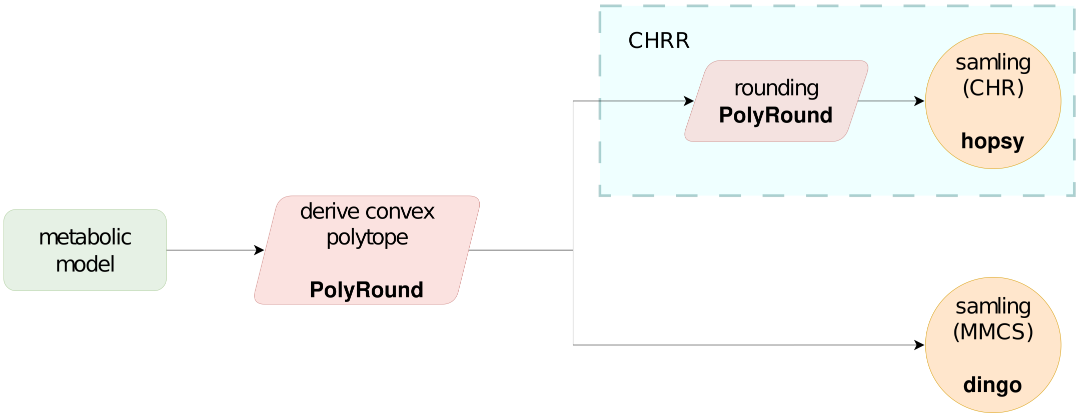


Figure 2: Workflow followed for comparing `dingo` with state-of-the-art sampling approaches. CHR: Coordinate Hit-and-Run; MMCS: Multiphase Monte Carlo Sampling.

We perform our benchmarks by requesting ESS $= 1000$ and PSRF $\leq 1.1$ for `dingo`. Since `hopsy` does not have an option to request a certain value of ESS we count its run-time until the generated sample achieves the same targets for both ESS and PSRF as in `dingo`. Last, in `hopsy` we use 5 chains following the same process as in [6].

Our benchmark dataset constitutes of five models from the BiGG Models knowledgebase,

[3]`https://github.com/modsim/hopsy`

Recon2 (version 2.2) from the BioModels database and Recon3D from Virtual Metabolic Human database.

Notably, `dingo`'s performance is comparable (i.e., same order of magnitude) to `PolyRound` that computes only the rounding. When comparing `dingo` to `PolyRound` for rounding plus `hopsy` for sampling, `dingo` is faster 2-13 times depending on the dimension. Last but not least, `dingo` can sample from recon3D and obtain an ESS = 1000 in around a day, while `hopsy` could not reach that ESS in 10 days. For details see Table 2.

For the reproducibility of our benchmark results note that we have used `dingo` version 0.1.0[4], `PolyRound` version 0.2.0 and `hopsy` version v0.2.0[5]. Also the scripts that we have used along with their outcomes (processed polytopes and samples) are publicly available under the following Zenodo repository:

https://doi.org/10.5281/zenodo.7581683.

#### 3 Illustrations and statistical tools

`dingo` provides a few illustrations and statistical tools that can help the user to make inference on a metabolic model. In particular, it provides probability density estimation for a flux distribution, given the marginal samples. Moreover, it provides copula estimation and plotting. A copula is a bivariate probability distribution for which the marginal probability distribution of each variable is uniform; it can be used to capture the dependency between two random variables (e.g., two reaction fluxes).

Recently, the virus biomass objective function (VBOF) of SARS-CoV-2 was generated and integrated with a genome-scale metabolic model of human alveolar macrophages by Renz *et al.* [17]. In their study, the authors performed Flux Balance Analysis (FBA) combined with reactions knock-out to reveal that the knock-out of Guanlyate Kinase 1 (GK1) decreased the growth of the virus to zero, while not affecting the one of the host. Among its several approaches [18], flux sampling can be used to increase the confidence level of FBA predictions [15]. As a demonstration, we performed flux sampling with `dingo` using the integrated human-virus model.

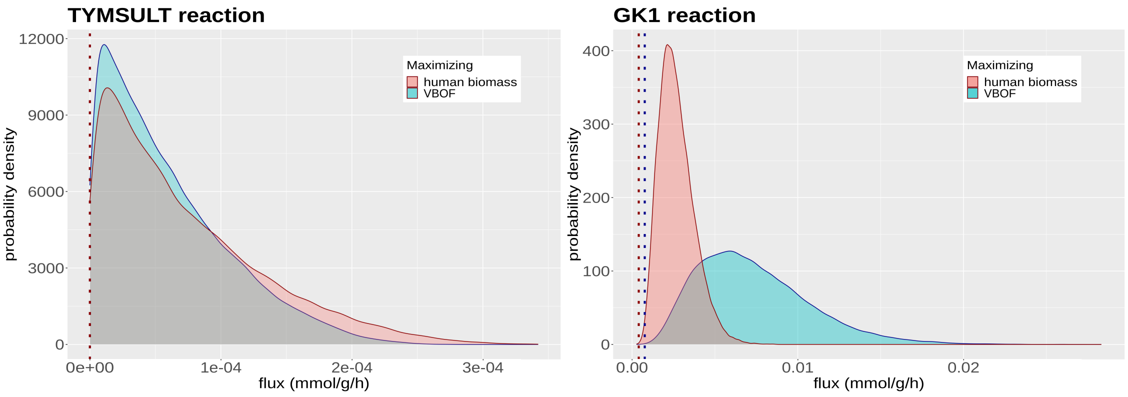


(a) (b)

Figure 3: Distributions of the integrated human-virus model reactions' flux values after maximizing for the host's biomass function (blue) and for the VBOF (red). (a) In case of the TYMSULT reaction, the distribution is the same for both cases (b) Contrary, the flux value distribution of GK1 shifts when maximizing for VBOF.

The flux space of the model was first sampled in an unbiased way, i.e., without requiring any optimization function to be optimized. Sampling was also performed after maximizing the

[4] commit hash #03d6e65a1278753bd879ea24f6512e7e8b95af93
[5] commit hash #4ad62ade6aa841d37f35d24e68043b3581358ce6

host's biomass function and finally, after maximizing the VBOF. In each of these cases, a flux value distribution for each reaction of the model was obtained, instead of a single value as in the FBA case. The plots in Figure 3 illustrate the distributions of the flux values of the Tyramine Sulfotransferase (TYMSULT) and the Guanylate Kinase 1 (GK1) reactions after maximizing for the host's biomass function (Figure 3(a)) and for the VBOF (Figure 3(b)) while the vertical lines correspond to the flux values computed by the FBA method. In the first case, FBA returned a zero value for both cases while the distributions were almost identical (Figure 3(a)), implying that the different requirements of host and virus are not related with the TYMSULT reaction. That is the case for the vast majority of the reactions of the model. On the other hand, FBA highlighted that the flux of GK1 increases when VBOF is maximized. That is further highlighted by the flux densities of GK1 reaction where both the mean and the variance of the distribution were increased when VBOF was maximized (Figure 3(b)).

To capture the dependency between the virus's biomass and the flux of the GK1 reaction, a copula of the GK1 and the human biomass fluxes' distributions was computed maximizing first for VBOF. As shown in Figure 4, their negative dependency further support the findings of Renz *et al.*. Interestingly, the FBA flux values for GK1, were rather lower than it is more likely for them to be, both if it is the human biomass that has been maximized or VBOF.

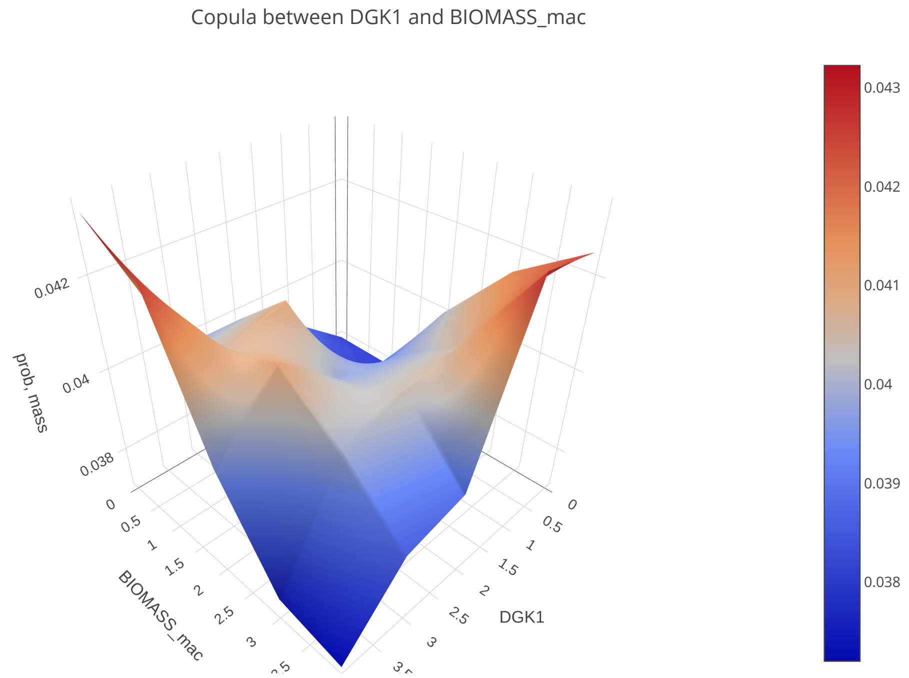


Figure 4: Copula of the biomass function and the GK1 reaction flux values' distributions, after maximizing the integrated model for VBOF. A positive dependency between the two reactions is shown as the flux of the virus biomass is low then most probably, the flux of GK1 is also low and the same applies in case of a high flux value.

In conclusion, flux sampling has the potential to be useful in drug targeting studies [16, 18].
